# Baseline cognitive abilities shape the effects of tDCS, tACS, and otDCS on memory

**DOI:** 10.64898/2026.02.25.707008

**Authors:** Jovana Bjekić, Marko Živanović, Carlo Minussi, Saša Filipović

## Abstract

Transcranial electrical stimulation (tES) can modulate neural dynamics, yet its effects on memory are heterogeneous. Individual differences in cognitive profiles, may well be one of the potential causes by setting boundary conditions on the extent and mode of the tES-induced modulation of network dynamics. In a sham-controlled, within-subject study (N = 42), we compared the effects of tDCS (1.5 mA), tACS (±1.0 mA at individual theta frequency, ITF), and otDCS (1.5 mA ± 0.5 mA at ITF) over the left posterior parietal cortex on object–location (OL) associative memory, and examined whether six cognitive abilities (figural reasoning, semantic, visuospatial, processing speed, working memory, mnemonic binding) moderate stimulation outcomes. Associative memory recognition improved selectively under theta-otDCS, whereas tDCS and tACS showed no significant group-level effects. Yet all tES protocols exhibited considerable interindividual variability. Relative to cognitive abilities, processing speed moderated tES effects in line with neural efficiency predictions, yielding greater gains in cognitively faster individuals. In contrast, mnemonic binding and figural reasoning moderated benefits in a compensatory manner, with larger improvements in lower-ability individuals. Overall, the effects of tES on associative memory were specific to the tES protocol and outcome measure while being strongly shaped by cognitive profile via complementing magnification and compensation mechanisms.

## Introduction

Associative memory (AM), the capacity to bind distinct elements of an experience into unified representations, is fundamental for everyday functioning. A relevant form of AM is object-location (OL) memory, our ability to remember and recall where objects belong in the environment (Postma et al., 2008). Early deficits in OL memory, such as frequently misplacing objects, are among the first cognitive markers of Alzheimer’s disease (Kessels et al., 2007; Postma et al., 2008). However, effective treatments remain lacking: pharmacological options and lifestyle interventions such as exercise, diet, or cognitive training offer only limited effects (Cummings et al., 2019; Hill et al., 2017; Martínez-López et al., 2024; O’Brien et al., 2017). Since memory relies on neural plasticity mechanisms (Kolb & Whishaw, 1998), it is reasonable to expect that memory performance should be susceptible to plasticity-based interventions such as non-invasive brain stimulation (NIBS; Filipović & Bjekić, 2025). Specifically, NIBS promote, via excitability changes, the neuronal reorganization phenomena that define plasticity, allowing these protocols to influence memory processes by modulating synaptic activity, and network-level dynamics. Accordingly, clarifying the mechanisms by which NIBS affect memory in healthy adults can optimize target engagement strategies, thereby increasing the likelihood of effective translation to clinical populations.

Due to its affordability, safety, and tolerability, transcranial electrical stimulation (tES) has emerged as a particularly attractive tool for memory-focused neuromodulation research. tES encompasses techniques that apply weak currents (typically 1–2 mA) to modulate cortical excitability balance and network dynamics, such as transcranial direct current stimulation (tDCS), transcranial alternating current stimulation (tACS), and oscillatory tDCS (otDCS). Anodal tDCS delivers constant positive-polarity currents thought to shift neuronal membrane potentials and bias cortical excitability (Filmer et al., 2014; Lefaucheur & Wendling, 2019), whereas tACS applies sinusoidal currents at defined frequencies to interact with and potentially entrain/modulate endogenous oscillations (Agboada et al., 2025; Antal & Paulus, 2013; Vogeti et al., 2022; Wischnewski et al., 2023). otDCS combines these principles by embedding oscillatory waveforms within a direct current offset, therefore aiming to induce synergic effects of baseline excitability changes by the direct component and targeted oscillatory entrainment (Guo et al., 2025; Vulić et al., 2021).

Although encouraging findings have been reported, the effects of tES on memory outcomes remain inconsistent (for review see Bjekić et al., 2023). Traditionally, this inconsistency has been attributed to methodological heterogeneity, mainly differences in the stimulation protocols, including stimulation parameters (e.g., intensity, duration, waveform type), and targeted brain regions (e.g., electrodes montage, electrical field distribution) (Vergallito et al., 2022). To address this, a range of tES-protocol optimization strategies have been introduced (for review see Van Hoornweder et al., 2025), including multielectrode montages to increase focality of tDCS (Fromm et al., 2024; Splittgerber et al., 2020), tACS (Lang et al., 2019), and most recently otDCS (Manojlović et al., 2025), as well as approaches to personalize frequencies (Živanović et al., 2022), or tailor current waveforms to cross-frequency coupling associated memory-relevant neural dynamics (Alekseichuk et al., 2016; Diedrich et al., 2024, 2025). Yet, even with these optimized protocols, outcome inconsistency on tES-memory studies remains substantial, underscoring the need to go beyond methodological and group-level effects and analyze inter-individual differences in tES responsiveness.

Cognitive functions emerge from coordinated interactions within distributed neural networks. Accordingly, enduring inter-individual differences in network efficiency and cognitive profile reflect functional states (e.g., excitability and connectivity), rendering tES effects on cognition inherently state dependent. Thus, baseline cognitive characteristics should act as boundary conditions that shape the extent and mode of stimulation effects, yet they remain insufficiently studied. Prior work suggests that the baseline level of memory capacity can constrain or facilitate responsiveness to anodal neuromodulation, such that individuals with lower initial performance sometimes benefit more from tES (Assecondi et al., 2022; Habich et al., 2017; Splittgerber et al., 2020), while those with higher abilities may show ceiling effects or even a decrease in performance (Gözenman & Berryhill, 2016; Splittgerber et al., 2020).

Therefore, baseline cognitive abilities can be particularly relevant for tES-induced effects on complex cognitive functions such as OL memory. Successful OL memory relies on distinct yet interacting operations: the processing and identification of an object, the assessment of its location (i.e., spatial processing), and the binding of these elements into an integrated representation (Postma et al., 2008). Performance on visuospatial OL tasks thus emerges from the joint contribution of multiple cognitive abilities. Namely, figural reasoning supports relational processing and the detection of relational patterns, semantic ability enriches encoding with conceptual knowledge, and visuospatial ability underpins the integration of objects with their spatial positions. Furthermore, processing speed facilitates efficient discrimination and visuospatial search, while working memory enables the maintenance and manipulation of recently encoded associations. Finally, mnemonic binding ability directly supports linking objects and locations into coherent representations. Individual differences in these abilities are expected to manifest not only in OL memory performance, such as that those with better binding abilities have higher performance, but also in the extent and mode of tES-induced changes in the task-relevant network functionality.

Hence, *individual differences in cognitive abilities* may shape responsiveness to tES via state-dependent, non-linear mechanisms, such that a person’s cognitive profile and the underlying excitation/inhibition balance of the task-relevant network(s) determine the extent and mode of stimulation effects. The baseline cognitive abilities set boundary conditions on the extent and mode of the tES-induced modulation of network dynamics (Miniussi & Bortoletto, 2025) and therefore, depending on the conditions, the same stimulation can either enhance or hinder performance. In this context, the baseline-dependency or compensation hypothesis predicts that stimulation effects vary inversely with baseline ability; stronger cognitive abilities leave less modulatory space (smaller gains), whereas weaker abilities may benefit when stimulation shifts the system toward a more functional regime. Conversely, according to the neural efficiency or magnification hypothesis, more efficient networks may show larger gains when external input aligns with and further optimizes already favorable states or engages synergistic nodes within the task-relevant circuitry. In practice, gains may depend on whether the stimulated network (potentially confined to specific subprocesses) adds functional leverage to the circuitry executing the task, thereby creating room for improvement. Therefore, baseline cognitive abilities can provide informative priors on brain function at the time of stimulation, helping to explain divergent constraining the extent and mode of tES-induced modulation outcomes across protocols and tasks.

In the present study, we systematically investigated the effects of three tES protocols - tDCS, tACS, and otDCS - on OL memory, assessed through both cued recall and recognition performance. We first evaluated and directly compared the group-level effects of the three tES protocols on distinct OL outcomes. In addition, we explored the effects on response times (RT) and analyzed within-group variability to understand if the tES effects on OL memory were heterogeneous or homogeneous across individuals. The central focus of the study was to examine the moderation of tES effects by individual differences in six cognitive abilities (figural reasoning, semantic ability, visuospatial ability, processing speed, working memory, mnemonic binding), and to compare these moderation patterns across tDCS, tACS, and otDCS. By integrating behavioral memory outcomes with inter-individual variability, the study sought to determine whether distinct tES protocols exert modality-specific effects on OL memory, and to elucidate how cognitive ability profiles shape the extent and mode of these effects.

## Methods

### Design

The study employed a fully repeated, sham-controlled, within-subjects cross-over design (Figure 1). In a pre-experiment, participants underwent an EEG session to estimate individual theta frequency (ITF) (Bjekić, Paunovic, et al., 2022), which parametrized the oscillatory stimulation protocols (tACS and otDCS). Prior to the experimental phase, participants completed a cognitive test battery to assess baseline cognitive abilities. Each participant then completed four experimental sessions, receiving anodal tDCS, tACS, otDCS, and sham stimulation in Latin-square counterbalanced order to control for order effects. Following each stimulation, participants completed a parallel version of the OL task, with versions counterbalanced across both sessions and conditions. The participants were blind to stimulation conditions, and two-experimenter procedure was implemented to maintain integrity of blinding. Sessions were spaced at least one week apart to minimize carryover effects.

**Figure 1.**
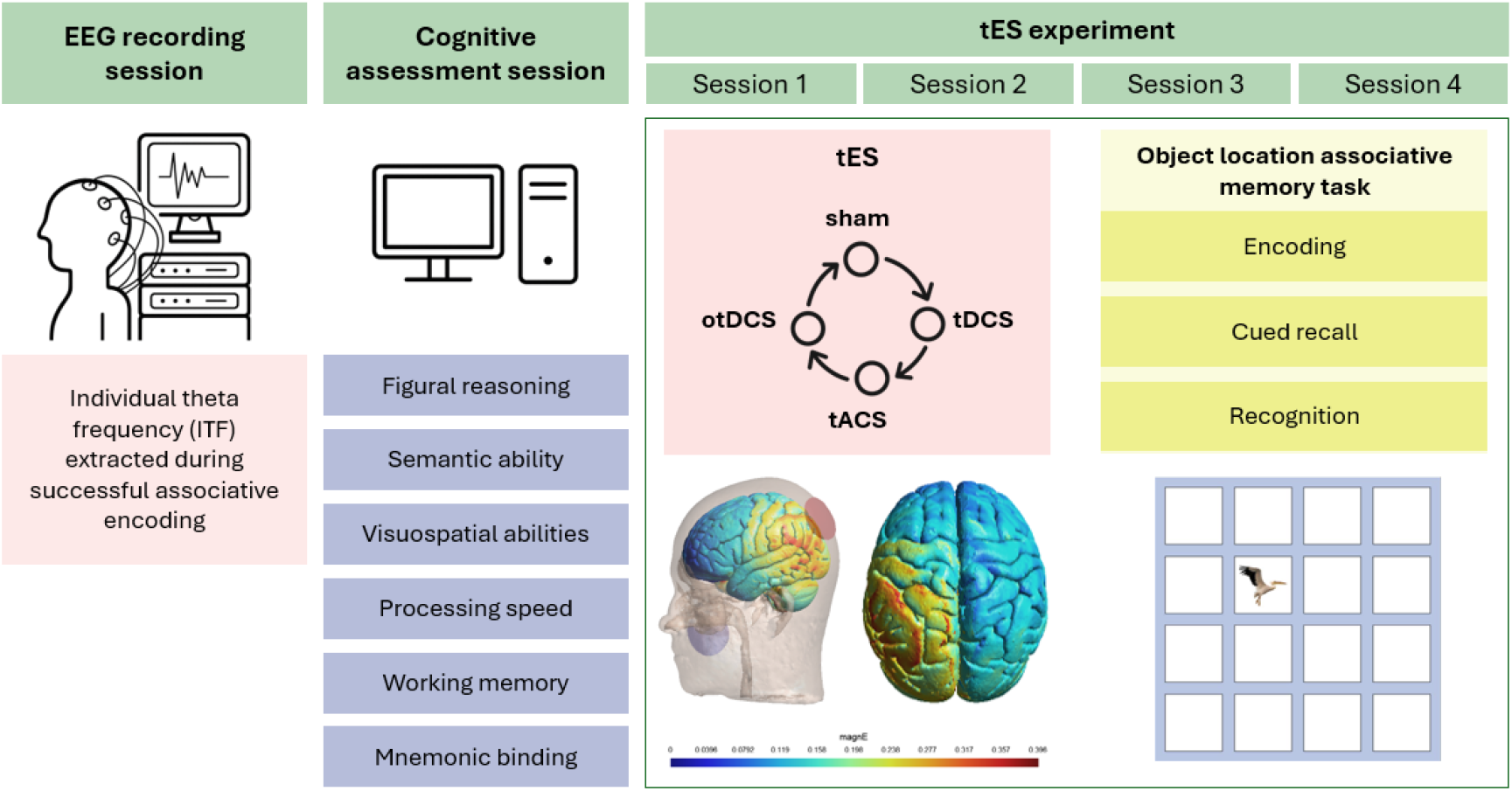
Overview of the experimental design and procedure. (1) EEG recording session, used to extract each participant’s individual theta frequency (ITF) during successful associative encoding, which was used in tACS and otDCS protocols; (2) Cognitive assessment session, in which six cognitive abilities were assessed; and (3) tES experiment, comprising four sessions (sham, tDCS, tACS, and otDCS) administered in a counterbalanced order. Following each tES, participants completed the object-location associative memory task with encoding, cued recall, and recognition block. Illustrations show stimulation montage (left PPC-cheek), modeled electric field distribution, and OL task layout.

### Participants

Forty-two healthy young adults (26 females) aged 22-34 years (*M* = 25.05, *SD* = 3.55) participated in the study. Inclusion criteria were right-handedness, normal or corrected-to-normal vision, and age between 20 and 35 years. Exclusion criteria were a history of seizures, neurological or psychiatric disorders, cognitive or learning disabilities, traumatic brain injury, chronic or acute dermatological conditions on the head, pregnancy or suspected pregnancy, and use of psychoactive substances or medication.

The study was approved by the Institutional Ethics Committee under the MEMORYST project (protocol code EO129/2020) and conducted following tES safety guidelines (Antal et at, 2017, Antal et al., 2025) and in accordance with the Declaration of Helsinki. All participants provided written informed consent and were compensated for participation.

### Transcranial electrical stimulation (tDCS, tACS, otDCS)

All tES protocols were delivered using NIC2 software-operated Starstim32 device (Neuroelectrics, Barcelona, Spain). Two 25 cm^2^ round rubber electrodes (≈ 5.6 cm diameter) inserted into saline-soaked sponge pockets (∼10 ml) were used, with impedance kept below 5 kΩ throughout stimulation. The target electrode (anode for tDCS) was positioned over the left posterior parietal cortex (PPC; P3 site of the 10-20 EEG system), a region supported by meta-analytic evidence as a key target for AM modulation in MRI-guided TMS studies (Badillo Goicoechea et al., 2025). The return electrode (cathode for tDCS) was placed on the contralateral cheek and secured with medical adhesive tape. This montage was successfully used in previous tES studies to modulate AM (Bjekić, Čolić, et al., 2019; Bjekić, Vulić, et al., 2019; Vulić et al., 2021; Živanović et al., 2022).

Each stimulation protocol lasted 20 min, including 30 s ramp-up (at the start), and ramp-down (at the end) periods. Four conditions were administered: (1) anodal tDCS, constant 1.5 mA current; (2) tACS, sinusoidal ±1 mA (2 mA peak-to-peak) at the ITF; (3) otDCS, sinusoidal ±0.5 mA (1 mA peak-to-peak) oscillating around constant 1.5 mA offset (range 1-2 mA) at the ITF; and (4) sham, a double-ramp up/down protocol with tDCS applied for 60 s (30 s ramp-up to 1.5 mA and 30 s ramp-down to 0 mA) at the start and at the end of the protocol to mimic tES-induced sensations (i.e., side effects) of active conditions. Anodal tDCS and otDCS were equivalent in terms of mean current intensity (1.5 mA), whereas otDCS and tACS shared oscillatory properties by being delivered at the ITF. The ITF ranged from 4.0-7.5 Hz (*M* = 5.18, *SD* = 1.17) and was determined using an off-line task-based dominant theta frequency extraction approach (Bjekić, Paunovic, et al., 2022).

### Associative memory task (object-location)

AM was assessed using the Animal-location task, an OL paradigm comprising three sequential blocks: Encoding, Cued recall, and Recognition. In the Encoding block, 32 animal images were presented sequentially for 2,000 ms each, located quasi-randomly within a 4 × 4 grid (16 possible locations). Participants were instructed to memorize the location of each animal. In the Cued-recall block, the probe animals were presented one at a time, and participants had to indicate the corresponding grid location using a mouse click. In the Recognition block, each animal was presented twice, once in its original (“old”) location and once in a re-paired (“new”) location, resulting in 64 recognition trials requiring an “old” versus “new” classification. Overall, the task lasted approximately 10 min.

Parallel task versions were used across stimulation sessions to minimize transfer effects. Versions differed in stimulus exemplars (e.g., different images of the cat/dog/small insect), background grid color, and spatial assignment of targets, which was randomized using a multiplication-by-substitution algorithm. The equivalence of task forms was established in prior validation study (Bjekić, Živanović, et al., 2022), demonstrating no systematic differences across versions for cued recall or recognition performance, with high inter-form equivalence (cued recall: *ICC* = 0.87, 95% CI [0.76 – 0.94]; recognition: *ICC* = 0.86, 95% CI [0.74 – 0.93]) and good internal consistency (cued recall: Cronbach’s α = 0.77 – 0.85; recognition: α = 0.76 – 0.80). The task was programmed in OpenSesame, materials and code are openly available via the Open Science Framework (OSF; https://osf.io/pmak3/).

From this task two main measures of AM were extracted: (1) **Cued recall success rate**: defined as the number of correctly recalled grid positions when participants were presented with a probe animal, in the cued-recall block; and (2) **Associative recognition accuracy**: quantified as ***d’***, a sensitivity index which is derived from hit rate i.e., correct identification of old pairings rate corrected for false alarms rate i.e., proportion of new pairings incorrectly identified as old [*d*′ = *Z*(hit rate) – *Z*(false alarm rate)]. In addition to *d*′, we also examined the **correct rejection success rate**, defined as the proportion of new pairings accurately identified as new. The latency for each response was recorded, and we calculated the average RT for cued recall (correct responses), hits (correctly identified target parings), and correct rejections (correctly classified new parings).

### Cognitive abilities assessment

A comprehensive battery of tests was administered to assess *individual differences in cognitive abilities* relevant to AM, comprising the tests detailed below.

a. **Figural reasoning**, i.e., the ability to identify patterns, rules, and logical relations among abstract visual stimuli, was assessed using the short form of Raven’s Progressive Matrices (sRPM). The test comprises 18 figural matrices, each missing the lower right element. The test requires inferring abstract rules from the presented figural elements and identifying the missing piece that logically completes the matrix among five options. Participants had 6 min to complete the task.
b. **Semantic abilities**, i.e., efficiency of accessing/retrieving meanings of the words from long-term memory, were assessed via the Synonym test (GSN) (Wolf et al., 1992). GSN is comprised of 39 items, each presenting a target word along with five possible answers. In each item, participants were required to select the word closest in meaning to the target word. The test had a 2 min time limit.
c. **Visuospatial abilities**, i.e., the capacity to analyze, integrate, and manipulate visual and spatial information, were assessed using the Spatial Imagination test (IT2) (Wolf et al., 1992). It consists of 39 items in which a target geometric figure was displayed alongside four unfolded cutouts. Participants were instructed to select the option that, when folded along the indicated lines, would form the target figure. They had 10 min to complete the test.
d. To assess **cognitive processing speed**, i.e., efficiency in identification and perceptual discrimination of simple objects, the Identical Figures test (IT1) was used (Wolf et al., 1992). In this test, participants were presented with 39 items showing a target image and four possible matches. Their task was to, as fast as possible, select the image that is identical to the target image. The time limit for this test was 4 min.
e. **Working memory** was assessed using the visual 3-back task (Živanović et al., 2021), in which participants are presented with a 3 × 3 grid on a white background where a black square appeared (1,000 ms) in a fixed interval (500 ms) in one of the nine quasi-random positions within the grid. Participants were instructed to respond only when the black square appeared at the same location as in three trials earlier. The task consisted of five blocks, each comprised of 32 stimuli. Each block lasted about 1 min, and the task’s total duration was about 5 min.
f. Finally, the **mnemonic binding** ability was measured with a nonverbal face-landscape AM task (Bjekić, Živanović, et al., 2022). In the encoding phase, participants studied 42 face-landscape pairs presented sequentially in fixed quasi-random order for 2000 ms each with variable interstimulus intervals (jittered between 1250-1750 ms). In the recognition phase, the 42 studied pairs were intermixed with 42 recombined pairs created by rearranging the same stimuli into novel pairings. Participants’ task was to judge each pair as “old” (studied) or “new” (new pairings). The participants took about 10 min to complete the test.

## Data Analysis

All analyses were conducted in JASP (Version 0.95.0) and R (Version 4.5.1). To evaluate the effects of tDCS, tACS, and otDCS on OL, we first examined group-level differences between active and sham stimulation using linear mixed-effects models (LMMs). Primary outcomes were accuracy measures (cued recall, recognition sensitivity d′, and correct rejection), whereas RTs were treated as secondary outcomes. Separate models were estimated for each stimulation modality and outcome. In all models, the contrast between active tES (tDCS/tACS/otDCS) and sham was entered as a fixed effect, session order was included as a covariate to account for repeated testing, and participant ID was included as a random intercept (e.g., *cued_recall_accuracy ∼ condition + session + (1* | *ID)*). Models were fitted using the lme4 (v1.1-37) and lmerTest packages (v3.1-3), and effect sizes are reported as partial eta-squared (η_p_^2^) alongside exact *p* values. Therefore, the model assessed if tES modulates memory performance, controlling for practice and interindividual variability. Next, to explore response variability we estimated residual variances for each condition and variance ratios relative to sham (e.g., *VR_otDCS =* σ*_otDCS /* σ*_sham*), providing insight into how dispersion varies across tES protocols beyond mean effects (see Supplementary for details).

To test whether interindividual differences on the level of cognitive abilities (figural reasoning, semantic ability, visuospatial ability, processing speed, working memory, mnemonic binding) moderated the effects of tES on AM outcomes (accuracy and RTs), the six cognitive measures (i.e., summary scores) were included as moderators in the extended prediction models *[e*.*g*., *cued_recognition_accuracy ∼ condition * (SRM + IT2 + IT1 + GSN + WM + AM) + session + (1* | *ID)]*. By entering all six cognitive measures simultaneously as moderators, the models estimated the unique variance explained by each cognitive measure while controlling for the overlap with the other moderators. Thus, the interaction term for a given cognitive ability reflects its specific moderating effect on the relationship between tES and AM outcomes, independent of shared variance with the remaining measures. Significant (*p* <.05) and trend-level (*p* <.10) condition × moderator interaction, were probed by estimating simple slopes of the tES effect at low (*M – 1 SD*), medium (*M*), and high (*M + 1 SD*) cognitive ability levels.

## Results

### Group-level effects on OL memory

Cued recall success remained comparable to sham across all tES conditions (tDCS: *F*_(1,39)_ = 0.010, *p* =.922; tACS: *F*_(1,39.068)_ = 0.004, *p* =.950; otDCS: *F*_(1,39.139)_ = 0.786, *p* =.381), though RTs, were slower after tDCS (*F*_(1,39)_ = 4.984, *p* =.031, ηp^2^ =.11) and otDCS (*F*_(1,39.115)_ = 5.329, *p* =.026, η_p_^2^ =.12).

Recognition accuracy (d’), did not differ from sham following either tDCS (*F*_(1,39)_ = 2.589, *p* =.116) or tACS (*F*_(1,39.153)_ = 0.050, *p* =.825). However, otDCS induced a significant improvement in associative recognition (*F*_(1,38.858)_ = 5.077, *p* =.030, η_p_^2^ =.12), indicating a selective enhancement in recognition sensitivity (Figure 2). No significant effects emerged specifically for correct rejection accuracy (tDCS: *F*_(1,39)_ = 1.150, *p* =.290; tACS: *F*_(1,39.098)_ = 0.189, *p* =.666; otDCS: *F*_(1,39.102)_ = 2.037, *p* =.161). No effects on correct recognition and correct rejection RTs were found either (Figure 2).

**Figure 2.**
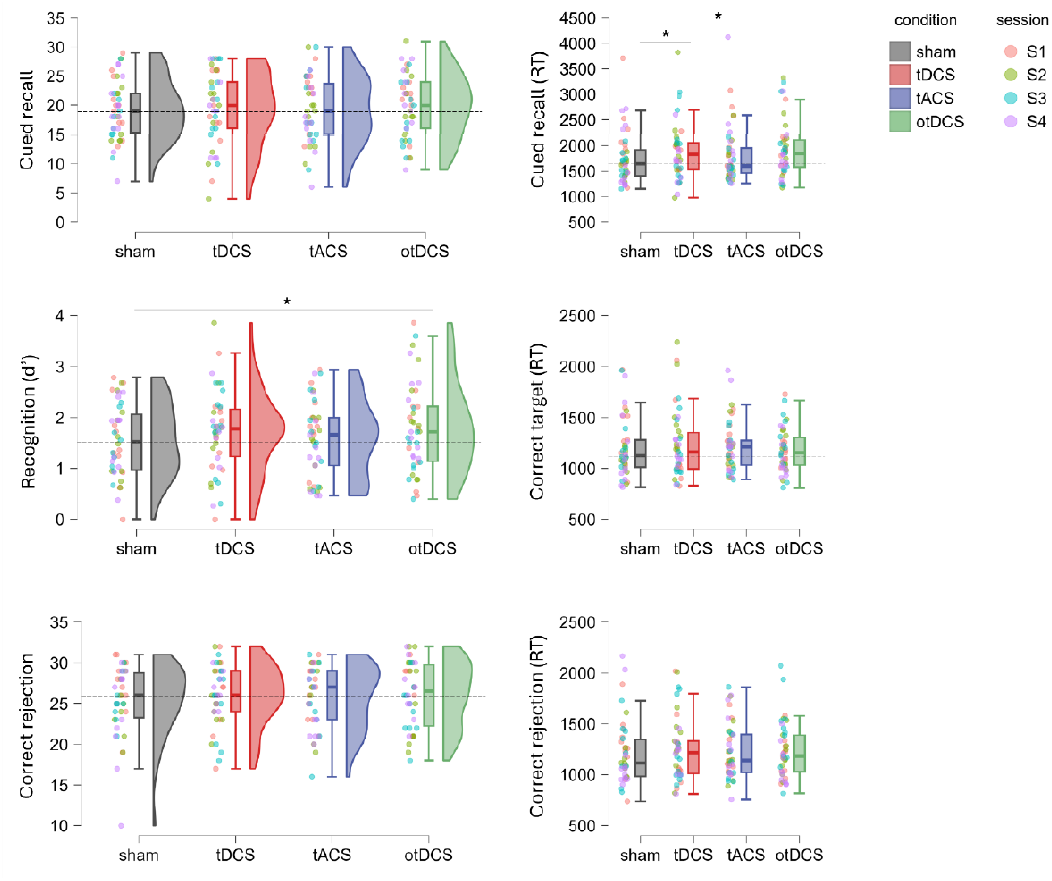
Group-level effects of sham, tDCS, tACS, and otDCS on object–location (OL) associative memory performance and response times (RT). Raincloud plots (left) depict OL memory outcomes: cued recall success rate (top), recognition sensitivity (d′; middle), and correct rejection success rate (bottom). Each plot combines a half-violin (distribution), boxplot (median and interquartile range), and individual data points, color-coded by session tES conditions are color marked: gray -sham, red - tDCS, blue – tACS, and green – otDCS. Box-plots (right) display RTs (in ms) for correctly recalled items (top), correctly identified targets – hits (middle), and correct rejections (bottom) across tES conditions. Significant differences (p <.05) are indicated with asterisks (*), and dashed lines denote the sham mean for visual reference.

Variability effects differed across tES protocols, consistently showing opposite patterns between accuracy and RT measures – the cued recall and correct rejection variability decreased at the same time when corresponding RTs increased, while the opposite was the case for correct recognition (Supplementary material). These bidirectional changes in variability were the most pronounced for otDCS.

### Moderation of tES effects by cognitive abilities

Out of six cognitive abilities analyzed, three (semantic abilities, visuospatial abilities, and working memory) did not show any moderator effect, while for other three various extents and modes of effects were found.

#### Cognitive processing speed

(i.e., efficiency/speed in identification and perceptual discrimination of objects) emerged as the most prominent moderator of tES effects across memory outcomes (Figure 3), with greater stimulation-related benefits observed in individuals with higher processing speed.

**Figure 3.**
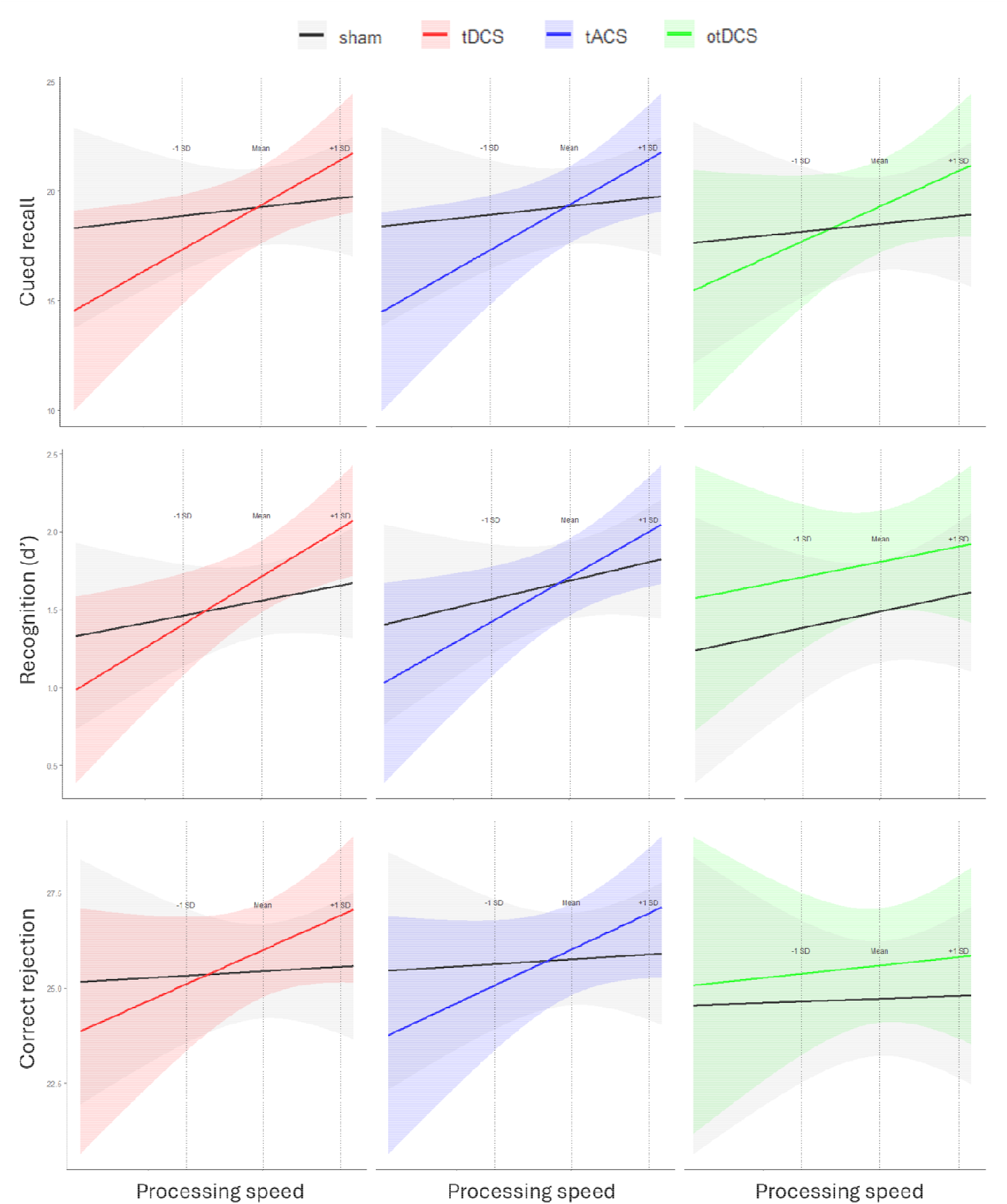
Cognitive moderator of tES effects on object location associative memory - Magnification pattern. Processing speed moderates the effects of tDCS (left), tACS (middle), and otDCS (right) on OL cued recall success rate (top), associative recognition sensitivity index - d′ (middle), and correct rejection success rate (bottom). Predicted OL scores (y-axes) are plotted as a function of processing speed (x-axes), vertical dashed lines denote processing speed levels at −1 SD (i.e., lower level of performance), the mean, and +1 SD (i.e., higher level of performance), black lines represent sham and tES protocols are color marked: red - tDCS, blue - tACS, and green - otDCS), with shaded areas indicating 95% confidence intervals.

Namely, for **tDCS**, processing speed moderated effects on all three measures – cued recall accuracy (*F*_(1,33)_ = 8.248, *p* =.007, η_p_^2^ =.20), recognition accuracy (*F*_(1,33)_ = 6.080, *p* =.019, η_p_^2^ =.16), and correct rejection accuracy (*F*_(1,33)_ = 4.282, *p* =.046, η_p_^2^ =.11). Specifically, individuals with higher IT1 scores improved their cued recall accuracy relative to sham (*t*_(33)_ = 2.265, *p* =.030), while those with lower IT1 scores declined (*t*_(33)_ = 2.122, *p* =.041). Similarly, higher processing speed was associated with greater improvements in recognition following tDCS compared to sham (*t*_(33)_ = 2.960, *p* =.006) and had higher performance gains as measured by correct reject accuracy (*t*_(33)_ = 2.317, *p* =.027).

The effects of **tACS** followed the same overall pattern, though they were slightly weaker. Processing speed significantly moderated cued recall accuracy (*F*_(1,32.922)_ = 5.375, *p* =.027, η_p_^2^ =.14) and correct rejection accuracy (*F*_(1,33.059)_ = 5.462, *p* =.026, η_p_^2^ =.14), while moderation of recognition accuracy was at trend level (*F*_(1,33.040)_ = 3.372, _p_ =.075, η_p_^2^ =.09). Higher IT1 scores were linked to a tendency for greater recall improvements following tACS (*t*_(33.1)_ = 1.819, *p* =.078), whereas lower IT1 scores were associated with a trend-level decline (*t*_(32.8)_ = 1.749, *p* =.090). Those with higher IT1 scores showed greater improvement in correct reject accuracy following tACS compared to sham (*t*_(33.1)_ = 2.091, *p* =.044), but the same tendency observed for overall recognition sensitivity did not reach statistical significance (*t*_(33.1)_ = 1.554, *p* =.130).

For **otDCS**, the moderating role of processing speed was less prominent. The effect of otDCS on cued recall accuracy was marginally moderated by processing speed (*F*_(1,33.056)_ = 4.144, *p* =.049, η_p_^2^ =.11), with higher IT1 scores predicting better recall performance (*t*_(33.1_) = 2.130, *p* =.041). However, unlike tDCS and tACS, processing speed did not moderate otDCS effects on recognition accuracy (*F*_(1,32.860)_ = 0.184, *p* =.670) or correct rejection accuracy (*F*_(1,32.917)_ = 1.353, *p* =.253).

#### Mnemonic binding

ability (i.e., nonverbal face-landscape AM task) also emerged as a significant moderator of tES effects, but selectively for correct rejection accuracy and showed a reverse moderation pattern to processing speed (Figure 4A). Namely, the effects on correct rejection accuracy were significantly moderated by mnemonic ability across all three stimulation conditions – tDCS *F*(_1,32.999)_ = 6.729, p =.014, η_p_^2^ =.17), tACS (*F*_(1,33.064)_ = 4.310, *p* =.046, η_p_^2^ =.12), and otDCS (*F*_(1,32.908)_ = 5.804, *p* =.022, η_p_^2^ =.15). In all cases, people with lower mnemonic binding ability tended to benefit more from stimulation. Specifically, significant stimulation-induced improvements in correct rejection accuracy were found for tDCS (*t*_(33)_ = 2.694, *p* =.011) and otDCS (*t*_(33.1)_ = 2.789, *p* =.009), while tACS followed the same pattern at a trend level (*t*_(33.1)_ = 1.868, *p* =.071).

**Figure 4.**
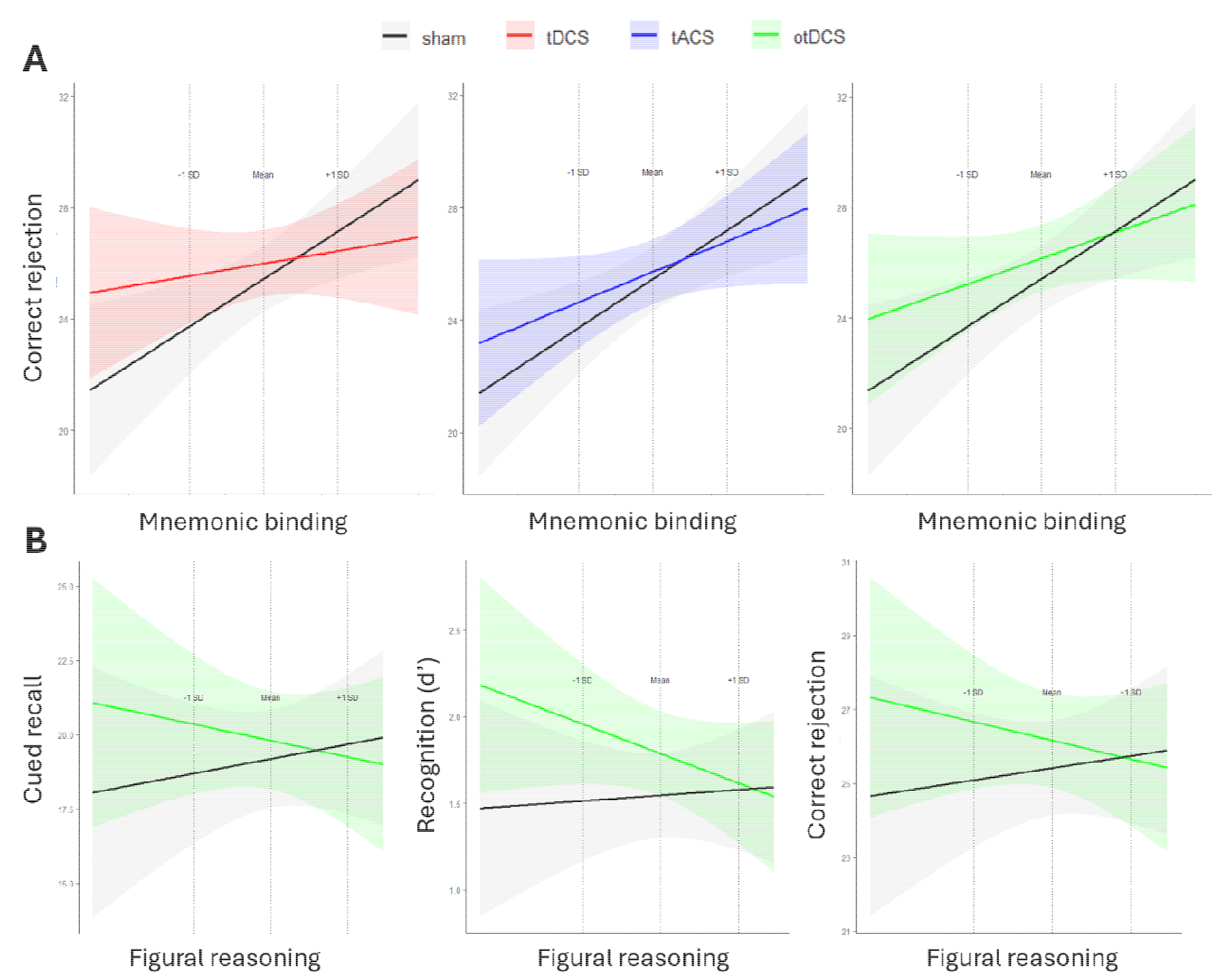
Cognitive moderators of tES effects on object location associative memory - Compensatory pattern. (**A**) mnemonic binding ability moderates the effects of tDCS, tACS and otDCS on correct rejection success rate. (**B**) Figural reasoning ability moderates otDCS effects across all OL measures (cued recall success rate, recognition sensitivity and correct rejection). Predicted OL scores (y-axes) are plotted as a function of cognitive ability (x-axes), vertical dashed lines denote cognitive ability levels at −1 SD, the mean, and +1 SD, black lines represent sham and tES protocols are color marked: red - tDCS, blue – tACS, and green - otDCS), with shaded areas indicating 95% confidence intervals.

#### Figural reasoning

(i.e., abstraction of rules, relations, and patterns on visual-abstract stimuli) selectively moderated the effects of otDCS on recognition performance (Figure 4B). Moderation analysis for recognition sensitivity (d’) (*F*_(1,32.830)_ = 2.866, *p* =.100, η_p_^2^ =.08) and correct rejection accuracy (*F*_(1,32.875)_ = 4.075, *p* =.052, η_p_^2^ =.11) revealed a consistent tendency toward greater stimulation-induced gains in lower-ability individuals. Specifically, recognition sensitivity improved significantly in participants with low figural reasoning (*t*_(32.9)_ = 2.563, *p* =.015), with a weaker but still significant effect in those with middle scores (*t*_(33)_ = 2.188, *p* =.036), whereas individuals with high figural reasoning ability showed no stimulation-related changes. The same pattern emerged for correct rejection accuracy, with greater improvements observed in participants with lower figural reasoning ability (*t*_(32.8)_ = 2.487, *p* =.018). The effects on cued recall descriptively mirrored the same pattern, which, however, did not reach statistical significance (*t*_(32.8)_ = 1.528, *p* =.136).

## Discussion

The present study systematically explored the effects of three tES protocols, tDCS, tACS, and otDCS, on OL memory by integrating group-level and individual differences analysis with pre-task baseline cognitive abilities assessment as potential source of variability of tES effects. Among stimulation protocols, only theta-otDCS improved memory performance at the group level, selectively enhancing associative recognition. Beyond this mean-level effect, interindividual variability was substantial, with processing speed, mnemonic binding ability, and figural reasoning reliably moderating stimulation outcomes. Taken together, the findings indicate that tES effects are both protocol- and process-specific, with memory performance emerging as the downstream outcome of an interaction between the individual’s baseline cognitive profile, the stimulation protocols, and the requirements of the memory task.

Improved OL performance following otDCS converges with previous evidence that theta-band otDCS can facilitate AM (Vulić et al., 2021; Živanović et al., 2022). By combining a tonic offset, which can induce a sustained network state, with rhythmic fluctuation, otDCS exerts both steady-state and oscillatory influences, biasing network excitability via subthreshold polarization and entraining intrinsic oscillatory activity, thereby potentially promoting hippocampal-cortical communication during memory retrieval. The selective enhancement of associative recognition suggests that otDCS made memory representations more accessible, but not sufficiently so that we could observe significant improvement in OL cued recall. This pattern accords with strength theory (Mickes et al., 2009; Wixted, 2007), as well as dual process models of memory in which recognition primarily reflects fast/automatic familiarity-based access, whereas recall depends on slower reconstructive retrieval (Buchler et al., 2008; Yonelinas, 2002). Furthermore, the results add to the body of literature showing heterogeneous tES effects across memory outcomes (Bjekić et al., 2023). This study further corroborates amassed evidence showing unreliable group level effects of tDCS and tACS on memory (Fromm et al., 2024; Klink et al., 2020). The variance and RT analysis, further highlight how mean effects obscure interindividual heterogeneity – what we typically observe as “mean net” effects is actually a result of tES inducing different levels of gains for different people. This heterogenization of performance levels, could be better understood by looking into the interaction between baseline cognitive profile and observed tES effects.

Processing speed emerged as the most robust predictor of stimulation benefits across memory outcomes. In line with the neural efficiency hypothesis, individuals with higher processing speed consistently benefited from tES, particularly tDCS and tACS, showing improved recall and recognition. In contrast, mnemonic binding displayed the compensatory pattern, with greater stimulation-induced gains among lower-ability individuals, particularly for tDCS and otDCS. These two abilities appear to have complementary roles: processing speed supports cognitive efficiency, reflected in faster information processing and more stable decision dynamics, whereas mnemonic binding governs the quality of associative links, enhancing selectivity and reducing false alarms. Both, however, reinforce the decision-useful signal to support AM performance, and can be viewed as indicators of the system’s functional state: processing speed reflects general neural efficiency, whereas mnemonic binding captures associative integrity. Accordingly, tES acts through two distinct yet convergent mechanisms, at the same time revealing how these abilities support AM through distinct mechanistic routes. Faster processors show magnification patterns, more efficient networks have higher cognitive processing speed; tES adds input consistent with this state and augments existing neural efficiency. Conversely, in low mnemonic-binding individuals, we observe compensatory effects, where tES promotes network-relevant activity to reduce ambiguity between similar associations (fewer false alarms, improved correct rejections), which, in the case of otDCS, leads to overall enhanced associative discrimination. Together, these complementary baseline-dependency mechanisms explain why identical stimulation protocols can produce divergent behavioral outcomes across individuals.

The differential susceptibility to tES is observed as a function of interaction between baseline cognitive abilities (which could be understood as prevailing functional state of the cognitive system) and tES modality (i.e., their different modes of action). Namely, processing speed moderated effects primarily under tDCS and tACS, consistent with their reliance on global excitability shifts and phase-dependent entrainment, both of which depend on existing processing efficiency. Baseline memory abilities seem to act as a boundary condition, with the tES-modality varying threshold for the beneficial effects. Figural reasoning effects were unique to otDCS, which, due to its asymmetric waveform, likely produces more complex subthreshold modulation that transcends the simple additive effects of tDCS and tACS, allowing individuals with weaker functional state to benefit.

Mechanistically, theta-otDCS appears to stabilize temporal dynamics of successful recognition consistent with theta-mediated coordination of hippocampal-cortical retrieval (Herweg et al 2020), thereby improving recognition accuracy. Unlike tDCS and tACS, its effects seem to be less dependent on baseline processing speed, likely because the combination of tonic and oscillatory components induces a sustained, more uniform network state across individuals. For cued recall, otDCS induced longer, yet more variable, recollection RTs without improving accuracy, suggesting that otDCS prompts individually tuned reweighting of the speed-accuracy trade-off, resulting in similar outcomes but via different temporal strategies. As with other tES protocols, effects are constrained by associative binding ability; however, otDCS adds a unique compensatory benefit for individuals with lower figural reasoning, providing an external temporal scaffold that promotes large-scale neural synchronization to facilitate information flow and enhances timing precision in the hippocampal-cortical communication circuit. By contrast, for high figural reasoning ability (i.e., networks already well-tuned), additional “optimization” by otDCS is redundant and may only alter retrieval strategy. Therefore, otDCS efficiency is determined by the degree of mismatch between the “quality” of endogenous dynamic and the task requirements. When the mismatch is larger otDCS provides “corrective” input but adds little or nothing when the functional state is already optimal.

From a practical standpoint, these findings underscore the importance of an individualized approach to cognitive neuromodulation, offering a clear path toward identifying cognitive abilities that predict responsiveness and tailoring tES protocols to individual functional profiles.

Methodologically, this study moves beyond mean-level analyses by integrating variance-based and moderation approaches, thereby avoiding issues inherent to previous baseline-dependency studies (Lega et al., 2022) and adopting an integrative framework across different tES modalities and memory outcomes to capture the complexity of tES-cognition interactions. Still, for a full mechanistic understanding, future studies combining behavioral and neuroimaging measures are needed, as behavioral data alone allow only indirect inference about underlying neural processes. Furthermore, the balance between magnification and compensatory mechanisms may differ in memory-deficient populations such as older or clinical groups. It is plausible that a certain level of cognitive efficiency is necessary for magnification mechanisms to operate, whereas compensatory effects may depend on the integrity of broader cognitive functioning.

## Conclusion

Together, these findings point out that the impact of tES on cognition cannot be fully understood without considering individual ability profiles. The same stimulation protocol may enhance performance in some individuals while impairing it in others, depending on whether stimulation amplifies efficient or invokes compensatory neural mechanisms. The combined evidence supports two complementary routes by which tES shapes OL memory: a magnification route, evident when networks are already well-performing (higher processing speed), and a compensation route, evident when relational/binding mechanisms are weak. These routes are not mutually exclusive and likely depend on the stimulated network, the process engaged (spatial recall vs. associative discrimination vs. mismatch detection), and the prevailing brain state, in line with state-dependent, non-linear models of stimulation (e.g., excitation/inhibition balance of the task-relevant network). (Miniussi & Bortoletto, 2025). Within this framework, individualized-theta otDCS appears especially suited to reorganize retrieval dynamics in ways that stabilize recognition timing and enable compensatory gains in discrimination for lower-ability profiles, while tDCS/tACS tend to amplify advantages in faster processors. Practically, reading tES effects through magnification and compensation clarifies why mean effects are modest, why variability is informative, and why tailoring protocols to cognitive profile and targeted subprocess may be necessary for understanding how tES modulates memory and obtain a robust memory enhancement.

## Acknowledgement

The authors would like to thank Dunja Paunovic, Katarina Vulić, Marija Stanković and Uroš Konstantinović for their role OL tasks development and data collection.

## Funding

This work was supported by EU-funded HORIZON Collaboration and Support Action TWINNIBS (101059369). MŽ and JB receive institutional support from the Ministry of Science, Technological Development and Innovation of the Republic of Serbia (451-03-137/2025-03/200163; 451-03-136/2025-03/200015). The funding body had no role in the study design, analysis and interpretation of data, writing of the report, and decision to submit the article for publication.

## Declaration of Competing interest

Authors declare that they have no competing financial or non-financial interests related to this work.

## Notes

### Competing Interest Statement

The authors have declared no competing interest.

