## Supplement for "Baseline cognitive abilities shape the effects of tDCS, tACS, and otDCS on memory"

Supplement 1

Variability analysis

**Analytic approach**: To explore whether response variability differed across stimulation conditions (sham, tDCS, tACS, otDCS), we compared homoskedastic and heteroskedastic models using the nlme package (v3.1-166). In heteroskedastic models, condition-specific residual variances were specified via a variance identity function [*weights = varIdent(~ 1 | condition)*]. Given that the study was not powered to detect small-to-medium effects on residual variance when compared using likelihood-ratio tests under REML estimation, results are primarily reported descriptively (may not generalize beyond the present sample), including estimated residual variances for each condition and variance ratios relative to sham (e.g., *VR_otDCS = σ_otDCS / σ_sham*). This approach allows to quantify how dispersion varies across tES protocols beyond mean effects.

**Results**:

For cued recall measures, in comparison to sham, the variability in cued recall success rate was unchanged under tDCS (*VR* = 0.987), but reduced following both tACS (*VR* = 0.643; ~36% lower) and otDCS (*VR* = 0.637; ~36% lower). In contrast, variability of RT for correctly recalled items increased across all stimulation protocols, most prominently under otDCS (*VR* = 2.447; ~145% higher than sham), followed by tACS (*VR* = 1.930; ~93% higher) and tDCS (*VR* = 1.652; ~65% higher). Therefore, it seems that tES tends to stabilize (homogenize) cued recall success rates across participants, but allows RTs to be more dispersed, suggesting diversification among participants in timing strategies.

For associative recognition measures, variability of sensitivity index d′ increased under all stimulation conditions relative to sham, with the largest increase after otDCS (*VR* = 1.707; ~71% higher), followed by tACS (*VR* = 1.423; ~42% higher) and tDCS (*VR* = 1.381; ~38% higher). By contrast, variability of recognition RTs for hits decreased under oscillatory protocols, particularly otDCS (*VR* = 0.395; ~60% lower) and tACS (*VR* = 0.519; ~48% lower), while remaining stable under tDCS (*VR* = 1.014). Therefore, tES, particularly otDCS stabilizes temporal dynamics of recognition across participants which leads to more variable success rates.

For correct rejection measures, variability in correct rejection success rate was reduced under tDCS (VR = 0.665; ~33% lower) and otDCS (VR = 0.661; ~34% lower), but unchanged under tACS (VR = 1.005). The variability of RTs for correctly rejected trials increased under tDCS (VR = 1.326; ~33% higher) and otDCS (VR = 1.321; ~32% higher), while remaining approximately the same after tACS (VR = 0.908), indicating that tES tends to uniform accuracy against more variable RTs.

**Conclusion:** tES does not affect performance level uniformly. tES, particularly otDCS it changes within-group variability. Here it appears that tES systematically reweights the speed-accuracy balance.
